# Distinguishing evidence accumulation from choice confidence in neural signatures of perceptual decision making

**DOI:** 10.64898/2026.10.29.760356

**Authors:** Daria Monakhovych, John P. Grogan, Simon P. Kelly, Redmond G. O’Connell

**Affiliations:** School of Psychology, Trinity College Dublin, Dublin, Ireland. Trinity College Institute of Neuroscience, Trinity College Dublin, Dublin, Ireland; School of Electrical and Electronic Engineering and UCD Centre for Biomedical Engineering, University College Dublin, Dublin, Ireland

**Keywords:** perceptual decision-making, confidence, prior knowledge, Electroencephalography (EEG), Centroparietal Positivity (CPP), mu-beta oscillations

## Abstract

Human electrophysiology research has identified the Centro-parietal Positivity (CPP) as an apparent neural correlate of sensory evidence accumulation during perceptual decision-making. The CPP has also been shown to closely scale with subjective confidence. Coupled with the fact of its domain-generality and lack of choice selectivity, this observation raises the possibility that the CPP might in fact trace subjective confidence rather than cumulative evidence, two quantities that are closely correlated in most experiments. To disentangle confidence from cumulative evidence, we used predictive cues with 80% validity to manipulate participants’ prior expectations during a contrast discrimination task while recording electroencephalography (EEG). At the time of decision termination, confidence is expected to reach a higher level for validly cued versus invalidly cued choices, whereas the opposite would be expected of a cumulative evidence signal due to biased decision starting points. As expected, valid cues led to faster, more accurate and more confident responses compared to invalid and neutral cues. At the neural level, cues produced systematic increases in preparation of the expected action prior to evidence onset as indexed by premotor mu/beta activity, consistent with cue-biased criterion adjustments. The CPP’s pre-response amplitude was significantly reduced for validly cued choices and the effect remained even when matching for confidence. Together these findings suggest that the CPP is not a direct read-out of subjective confidence, but an evidence accumulation signal that contributes to both decision formation and confidence estimation.

## Introduction

Neurophysiological studies across a range of species have identified brain signals that trace the process of perceptual decision making as it unfolds (Carandini & Churchland, 2013; Gold & Shadlen, 2007; Hanks & Summerfield, 2017; Kiani et al., 2008; O’Connell & Kelly, 2021). These decision signals exhibit key dynamical properties of sensory evidence accumulation that conform to the predictions of the dominant mathematical decision models including a gradual evidence-dependent, response time-predictive build-up to a peak immediately prior to response (Kelly & O’Connell, 2013; Loughnane et al., 2016; O’Connell et al., 2012; Steinemann et al., 2018). The signals that have been identified appear to fall into distinct functional categories. Most are ‘effector-selective’, tracing the preparation of a specific decision-reporting action (de Lange et al., 2013; Donner et al., 2009; Steinemann et al., 2018; Twomey et al., 2016). In addition, a distinct class of decision signal has been observed in the human electroencephalography (EEG) which exhibits dynamics consistent with evidence accumulation irrespective of the task’s action requirements and even when no overt action is required at all to report choices (Kohl et al., 2020; O’Connell & Kelly, 2021; van Vugt et al., 2019).

Aside from its movement-independence, the Centro-parietal Positivity (CPP) exhibits additional functional differences from effector-selective decision signals in the human EEG. Effector-selective signals such as lateralised mu/beta (MB) activity show strategic starting-level adjustments in response to experimental manipulations of time pressure or prior knowledge and reach a stereotyped amplitude over contralateral sites immediately prior to the choice report consistent with a threshold-crossing effect (Kelly et al., 2021; Steinemann et al., 2018; Walsh et al., 2024). In contrast, the CPP does not show such starting-level changes, while its final pre-choice amplitude systematically varies as a function of time pressure, prior knowledge, reaction time (RT) and accuracy (Kelly et al., 2021; Kohl et al 2020; O’Connell et al., 2012; Steineman et al., 2018).

Simulations derived from sequential sampling models (e.g. Kelly et al., 2021) confirm that these characteristics are consistent with the CPP tracking a sensory evidence accumulation process whose duration and extent is determined by strategic adjustments implemented in down-stream motor areas.

However, another possibility that has been highlighted is that the CPP may track the observer’s subjective confidence in their choice (Urai & Pfeffer, 2014). Consistent with this idea, the CPP is positive in polarity irrespective of which alternative the participant chooses and its pre-response amplitude has been shown to scale with subjective confidence reported simultaneously with and subsequent to the initial choice (Dou et al., 2024; Feuerriegel et al., 2022; Gherman & Philiastides, 2015; Grogan et al., 2023, 2026; Herding et al., 2019). Given that subjective confidence and cumulative evidence are closely intertwined in most perceptual choice experiments (Hebart et al., 2016; Kiani et al., 2014; Murphy et al., 2015; Pleskac & Busemeyer, 2010), few previous studies have been able to definitively test these alternative functional accounts of the CPP. One recent study did show that it was possible to experimentally alter participants’ average confidence judgments without affecting their choice accuracy and without impacting on the amplitude of the CPP, a result that supports the evidence accumulation account (Desender et al., 2025).

Here we build on the work of Desender et al. (2025) to further establish whether the CPP relates most directly to evidence accumulation or subjective decision confidence. Participants performed a two-alternative contrast discrimination task in which they simultaneously reported their choice and their subjective confidence, following a cue which predicted the correct choice with 80% probability. Previous studies have shown that, under similar task conditions, the CPP’s pre-response amplitude is smaller on trials in which the correct alternative was validly cued compared to invalidly cued trials, consistent with cue-biased starting-level adjustments identified in behavioural model fits (Kelly et al., 2021; Kohl et al., 2020). However, these studies did not measure their participants’ subjective confidence. If the CPP reflects sensory evidence accumulation, its pre-response amplitude should be smaller for correct choices on validly compared invalidly cued trials that are matched for subjective confidence ratings. Alternatively, if the CPP reflects subjective decision confidence, its amplitude should not differ between these matched trials.

## Results

### Effects of Predictive Cues on Behaviour

We analysed behavioural and EEG data from 19 participants who performed a two-alternative forced-choice contrast discrimination task (Figure 1a). On each trial, participants judged which of two overlaid gratings appeared to have higher contrast and simultaneously rated their confidence in that decision. Choice and confidence were reported with a single left- or right-hand keypress (Figure 1a). We opted to have participants report their choices and confidence simultaneously to exclude the possibility that post-choice deliberation would complicate or obscure the relationship between pre-choice measurements of neural activity and confidence (see Grogan et al 2023; Grogan et al., 2026). At the beginning of each trial, a cue was presented that was either predictive of the correct alternative (with 80% accuracy) or non-predictive (‘neutral’ cue). Prior to commencing the experiment, task difficulty was titrated to ensure that participants’ accuracy on neutral trials was between 60-80% to avoid ceiling effects and to ensure substantial cross-trial variability in confidence levels.

**Figure 1.**
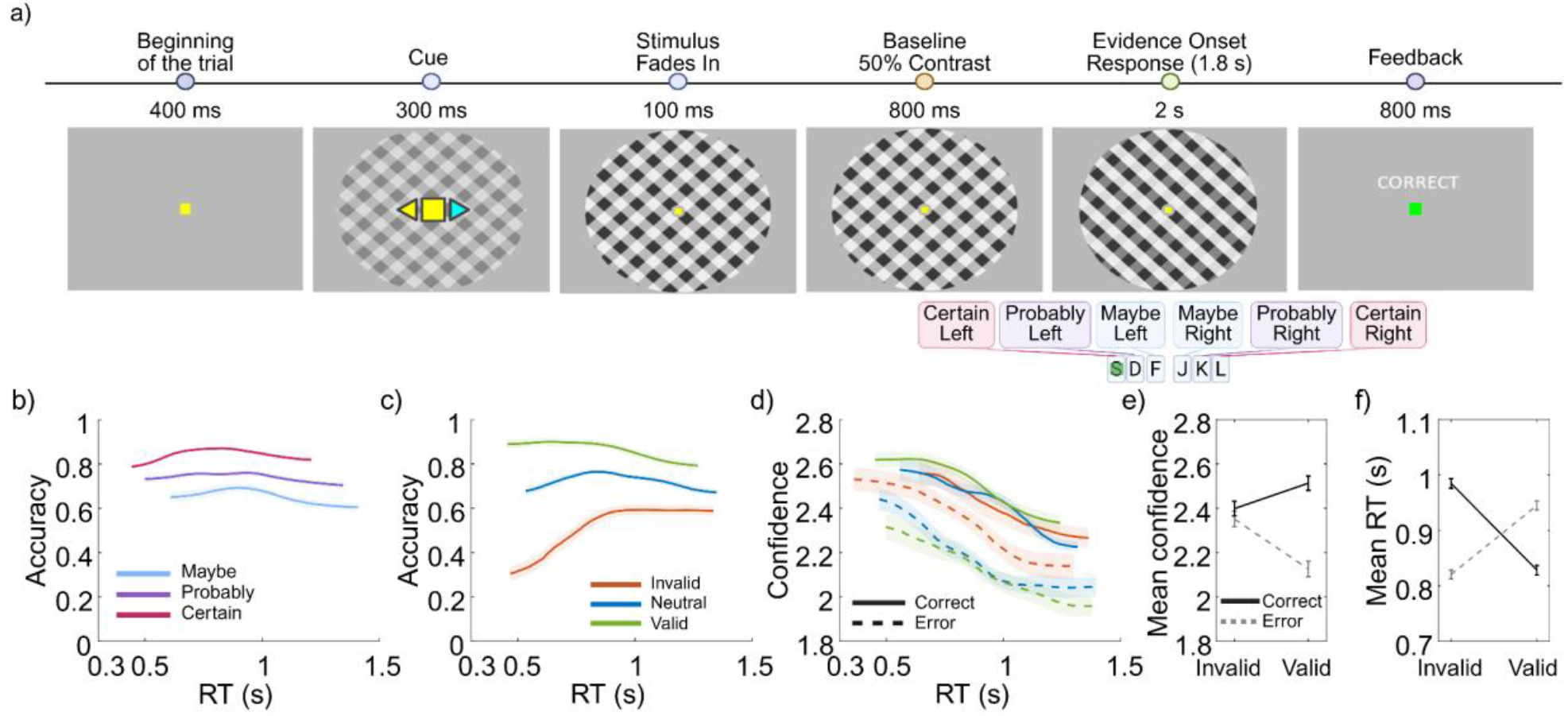
Task design and cue effect on behaviour. (a) Contrast discrimination task. Two overlaid gratings with the same contrast level tilted to the left and to the right faded in over 400ms during which time a predictive cue (coloured arrows) was presented either side of fixation. Following the fade-in, the gratings were then presented for an 800ms baseline interval to allow visual-evoked potentials to resolve in the EEG. Evidence onset took the form of an increase in the contrast of one grating and a decrease in the contrast of the other for 2 seconds, during which participants were required to report which grating was brightest and to simultaneously report their confidence in that judgment within a deadline or 1.8 seconds. Responses were made with a left- or right-hand button click, with S, D and F keys clicked using the left-hand ring, middle and index finger respectively corresponding to a certain, probably and maybe left choice and J, K and L clicked using the right-hand index, middle and ring finger respectively for maybe, probably and certain right choice. Feedback on their response was provided immediately after stimulus offset. The stimulus sizes on the figure are not scaled. (b-d) Conditional response-time functions calculated via sliding window analysis (20% RT windows, 1st-80th percentile; mean +-SEM across participants). (b) Accuracy as a function of RT across Confidence levels. Participants showed lower accuracy and slower RTs as their confidence decreased. (c) Accuracy as a function of RT for different cue types. Irrespective of RT, accuracy was highest on valid cue trials and lowest on invalid trials. RTs were fastest on valid trials and slowest on neutral ones. (d) Confidence as a function of reaction time, separated by cue types and accuracy. Overall confidence decreased rapidly with increasing RT. For correct trials, participants were fastest and most confident on valid trials, and slowest with lowest confidence on invalid trials, with neutral trials falling in between. The reverse pattern was observed for incorrect trials; participants were fastest and most confident on invalid cue trials (n=19). (e) Mean confidence was higher on correct than incorrect responses, increasing from invalid to valid cue trials for correct responses but decreasing for incorrect responses. (f) Mean RT on valid cue trials was quicker for correct than incorrect responses, whereas the opposite pattern was observed on invalid trials.

As is typically observed, Confidence levels were positively associated with choice accuracy at the trial-level (F (1,36131) = 816.19, β = 0.37, p < .001) and negatively associated with RT (F (1,36131) = 1794.5, β = –0.22, p < .001) (Figure 1b). Choice accuracy also varied significantly as a function of Cue Validity (F (2, 36130) = 1828.1, p < .001) with increased accuracy for valid compared to neutral cue trials (β = 0.97, p < .001) and for neutral compared to invalid cue trials (β = 0.95, p < .001). Similarly, RTs differed systematically by Cue Validity in the expected way (F (2, 36130) = 212.96, p < .001) with faster RTs following valid compared to invalid cues (β =-0.16, p < .001) and slower RTs following neutral compared to both valid (β = 0.26, p < .001) and invalid (β = 0.09, p = .001) cues (Figure 1c). Confidence was also modulated by Cue Validity (F (2, 36163) = 65.03, p < .001), being higher following valid compared to invalid cues (β = 0.096, p < .001) and reduced following neutral compared to both valid (β = - 0.14, p < 0.001) and invalid (β = -0.043, p = .012) cues.

We also observed a commonly reported Cue Validity x Correctness interaction (F (2, 36127) = 354.53, p < .001), driven by opposite effects of cue validity on correct versus incorrect RTs. That is, correct RTs were faster on valid trials (β = –0.29, p <.001) and slower on invalid trials (β = 0.14, p < .001) compared to neutral trials, while incorrect RTs were faster on invalidly cued trials compared to neutral trials (β = -0.3, p < .001) but did not significantly differ between valid and neutral trials (β = 0.056, p = .072) (Figure 1d, f).

A corresponding reversal of effects was also observed for confidence (Cue x Correctness interaction: F (2, 36127) = 354.53, p < .001). For correct responses, confidence was higher following valid compared to neutral cues (β = 0.11, p < .001), while there was no significant difference between neutral and invalid cue (β = 0.01, p = .72, Figure 1d, e). For incorrect responses, confidence was higher following invalid cues (β = 0.223, p < 0.001) and lower following valid cues (β = -0.11, p < 0.001) compared to neutral cues (Figure 1d, e).

### The Influence of Predictive Cues on Neural Decision Signals

Previous studies have shown that motor preparation signals exhibit systematic starting-level adjustments in response to predictive cues, with greater preparation of the cued response prior to sensory evidence onset (de Lange et al., 2013; Kelly et al., 2021). To verify that the predictive cues produced the expected strategic decision-process adjustments in the present study we first examined premotor MB lateralisation during the interval between the cue and evidence onset. For MB lateralisation, more negative values indicate relatively greater preparation of the cued response on cued trials and greater preparation of the chosen response on neutral cue trials.

Visual inspection of the data (Figure 2a) confirmed that there was strong cue-evoked lateralisation toward the cued hand prior to evidence onset. A mixed-effects model confirmed a significant main effect of Cue Validity(F (2, 36163) = 31.5, p < .001), with significantly greater lateralisation for valid cues (β = –0.11, p < .001) and invalid cues (β = –0.12, p < .001) compared to neutral, while there was no difference between valid and invalid trials (β = 0.01, p = .47).

**Figure 2.**
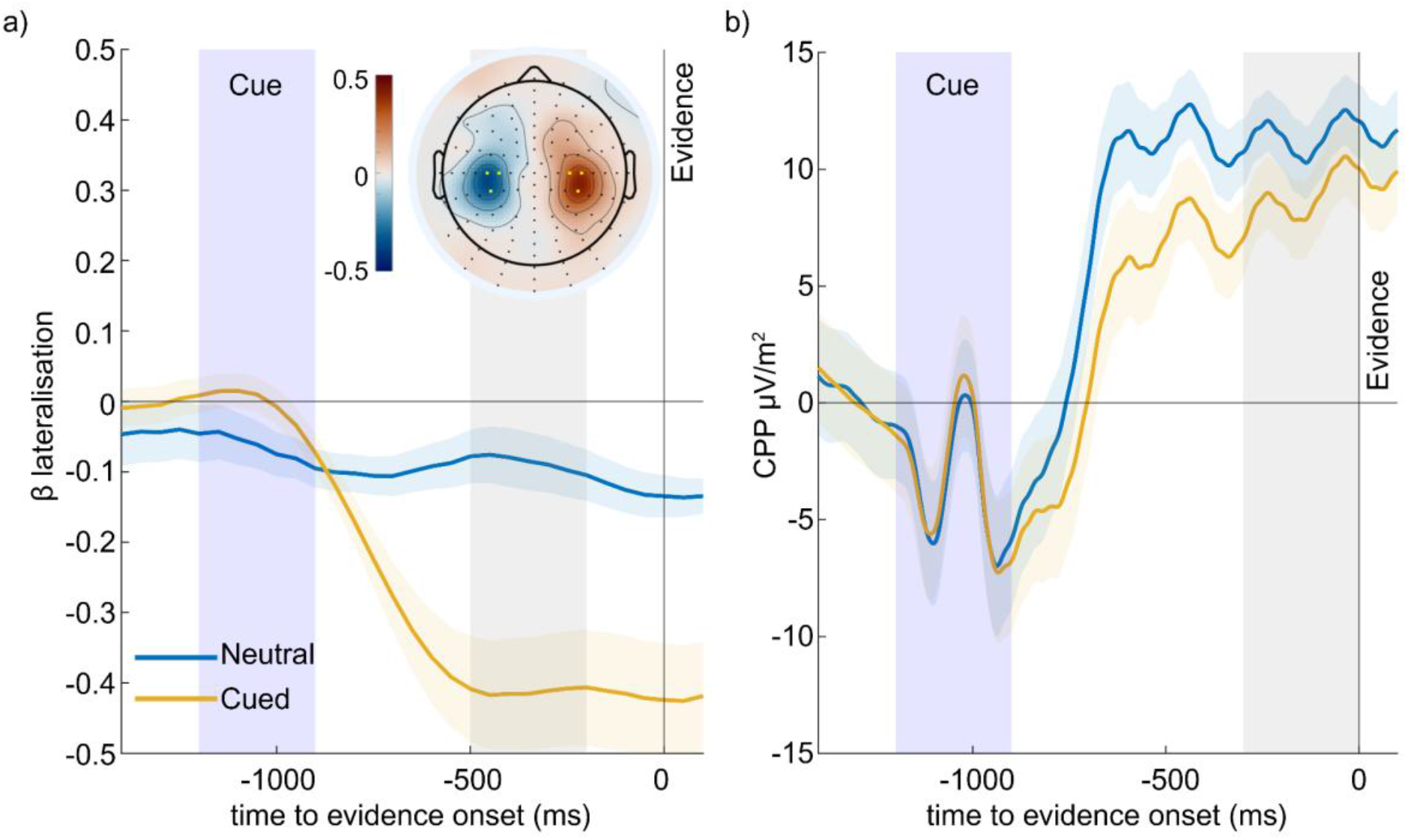
Effect of predictive cues on MB lateralisation and CPP amplitude prior to evidence onset when gratings were held at 50% contrast. (a) MB lateralisation (contralateral minus ipsilateral) was calculated relative to the direction indicated by the cue and eventual response on neutral cue trials. Strong MB lateralisation in the cued direction is observed on cued trials. (b) CPP waveforms across all trials and split by cue types show greater mean amplitude on neutral compared to cued trials. Blue shading (a, b) indicates the cue presentation period, while grey shading (a, b) shows pre-evidence time window used to calculate mean signal amplitudes for statistical analyses.

Turning to the CPP, we examined how its amplitude varied as a function of Cue Validity, and Correctness by fitting linear mixed-effects models including all main effects and their interactions for pre-evidence and pre-response CPP amplitudes separately.

Figure 2b highlights differences in the CPP’s amplitude between Neutral and Cued trials before the sensory evidence was presented. This was confirmed by a significant effect of Cue Validity on pre-evidence CPP amplitude (F (2,36121) = 4.52, p = .01) with larger amplitudes on neutral cue trials compared to invalid (β = -0.04, p = .035) and valid cues (β = -0.04, p = .002). These activity differences might reflect the fact that neutral cues were relatively rare and/or a greater tendency to initiate evidence accumulation prematurely in the absence of a cue (Devine et al. 2019). Since these amplitude differences occur during the interval used for applying baseline-correction to our post-evidence signal analyses (-150 to 0ms relative to evidence onset) and our hypotheses all centred on comparing of validly to invalidly cued choices (no difference in amplitude between these trials, β = 0.002, p = .91) we opted to remove the neutral trials from the remaining CPP analyses.

Pre-response CPP amplitude was significantly modulated by Cue validity (F (1,30150) = 12.67, p < .001). Despite participants reporting higher confidence on validly cued trials, pre-response CPP amplitudes were significantly smaller for validly compared to invalidly cued trials (β = -0.057). CPP amplitudes were also larger for correct compared to error responses (F (1,30150) = 16.22, β = 0.042, p < .001). In addition, whereas confidence was greater following invalid compared to valid cues on correct trials and following valid compared to invalid cues on error trials (Figure 3f), the CPP exhibited the opposite pattern although the Cue x Correctness interaction did not reach significance (F (1,30150) = 3.4, p = .065). A separate LMM including Confidence as a predictor showed that CPP amplitudes were modulated by Confidence (F (1,30146) = 9.65, β = 0.05, p =.001) as well as Cue Validity (F (1,30146) = 12.25, p=0.0004). The Confidence x Cue interaction did not reach significance (F (1,30146) = 3.3, p = .07), indicating that the positive relationship between CPP and confidence was preserved across cue types (Supplementary Figure 1).

**Figure 3.**
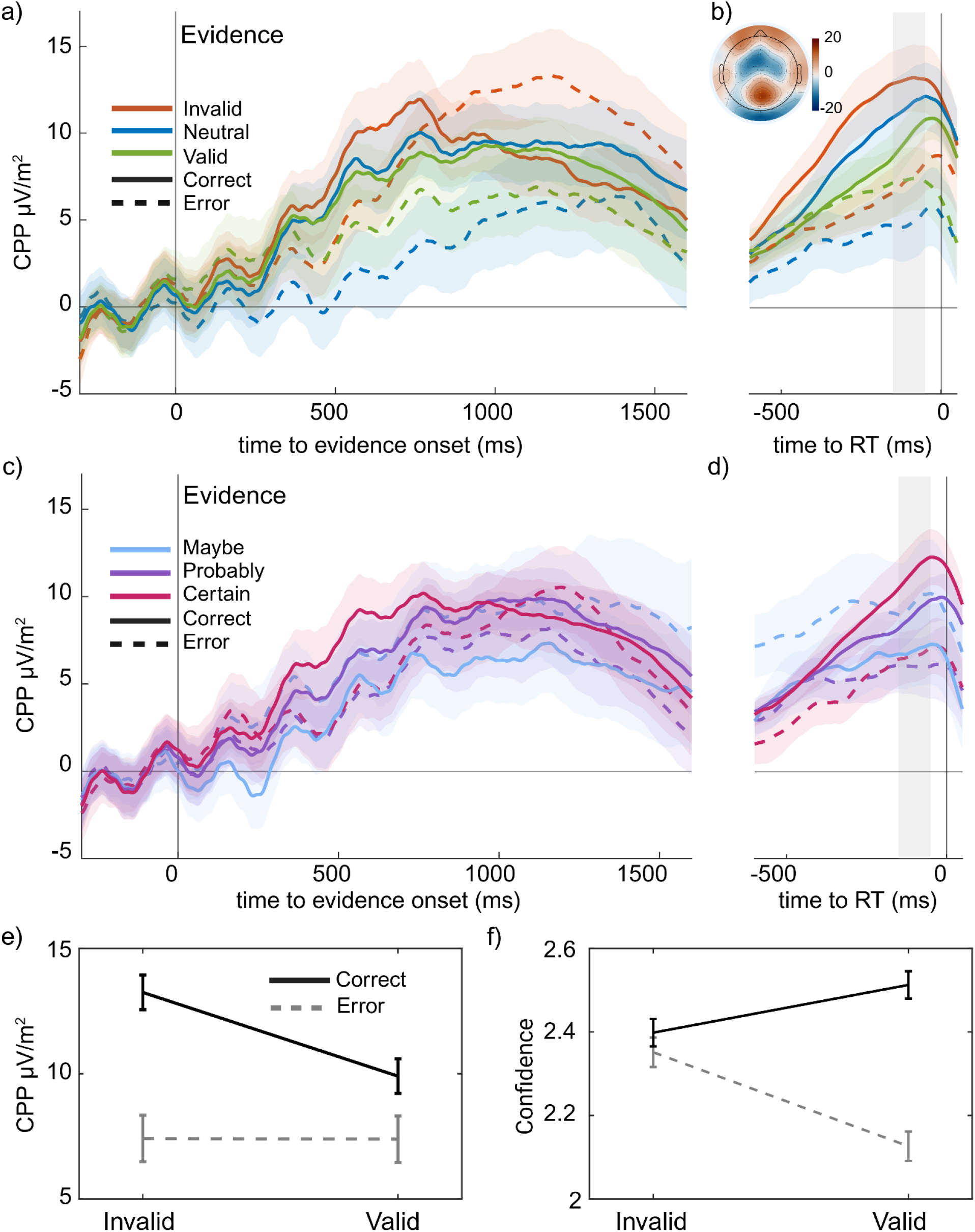
The CPP decision signal is modulated by predictive cues and scales with confidence. (a) Evidence-locked and (b) response-locked CPP split by cue types and correctness across all trials show lower mean pre-response amplitude for “valid” cue relative to “neutral” or “invalid” conditions on correct trials. (n = 19). (c) Evidence-locked and (d) response-locked CPP amplitudes plotted as a function of confidence and correctness across all trials. Pre-response CPP amplitudes were higher on trials where participants reported being ‘certain,’ compared to trials rated as ‘probably’ or ‘maybe’ (n = 19). (e, f) Mean pre-response CPP amplitudes (e) were smaller on validly compared to invalidly cued trials, even though confidence ratings (f) were higher (n = 19). Grey shading (b, d) shows the time window used for calculating mean pre-response CPP amplitudes for analyses. The inset plot (b) shows the grand-average topography for the –150 to –50 ms pre-response window for valid cue trials.

As there were substantial imbalances in trials numbers across our conditions (e.g. valid incorrect trials were rare) we sought further confirmation of these results by isolating trials in which participants made the correct choice and selected the ‘certain’ confidence rating and fit a separate statistical model which included the effects of Cue Validity, RT, and their interaction. Even when controlling for confidence in this way, Cue Validity was found to significantly influence pre-response CPP (F (1, 14709) = 14.2, p < .001), with significantly reduced amplitudes following valid cues compared to invalid cues (β = –0.098). There also was a significant Cue x RT interaction (F (1, 14709) = 7.71, p = .005), as pre-response CPP amplitude increased with RT on valid trials (β = 0.059, p < .001), while there was no significant relationship with RT on invalid trials (β = –0.025, p = .38).

## Discussion

In this study we investigated whether the CPP primarily reflects subjective confidence or sensory evidence accumulation during perceptual decision-making. We used predictive cues to manipulate participants’ expectations and hence produce opposing effects on the levels of confidence and cumulative evidence that would be reached by the time of choice commitment. As expected, the cues produced strong behavioural biases that were underpinned by corresponding biases in starting levels motor preparation prior to evidence onset, consistent with participants reducing the quantity of sensory evidence required for commitment to the expected response (de Lange et al., 2013; Kelly et al., 2021; Walsh et al., 2024). In addition, participants reported increased confidence on validly cued compared to invalidly cued trials (Tarasi et al., 2025). Given these observations, if the CPP were a signature of subjective confidence we predicted it should reach larger pre-response amplitudes on validly cued trials while if it were a signature of cumulative evidence, we predicted it should be larger on invalidly cued trials. The present data clearly supported the latter alternative with the CPP being significantly smaller on validly cued trials even when analyses controlled for RT and were restricted to correct responses matched for reported confidence (Fig.4).

**Figure 4.**
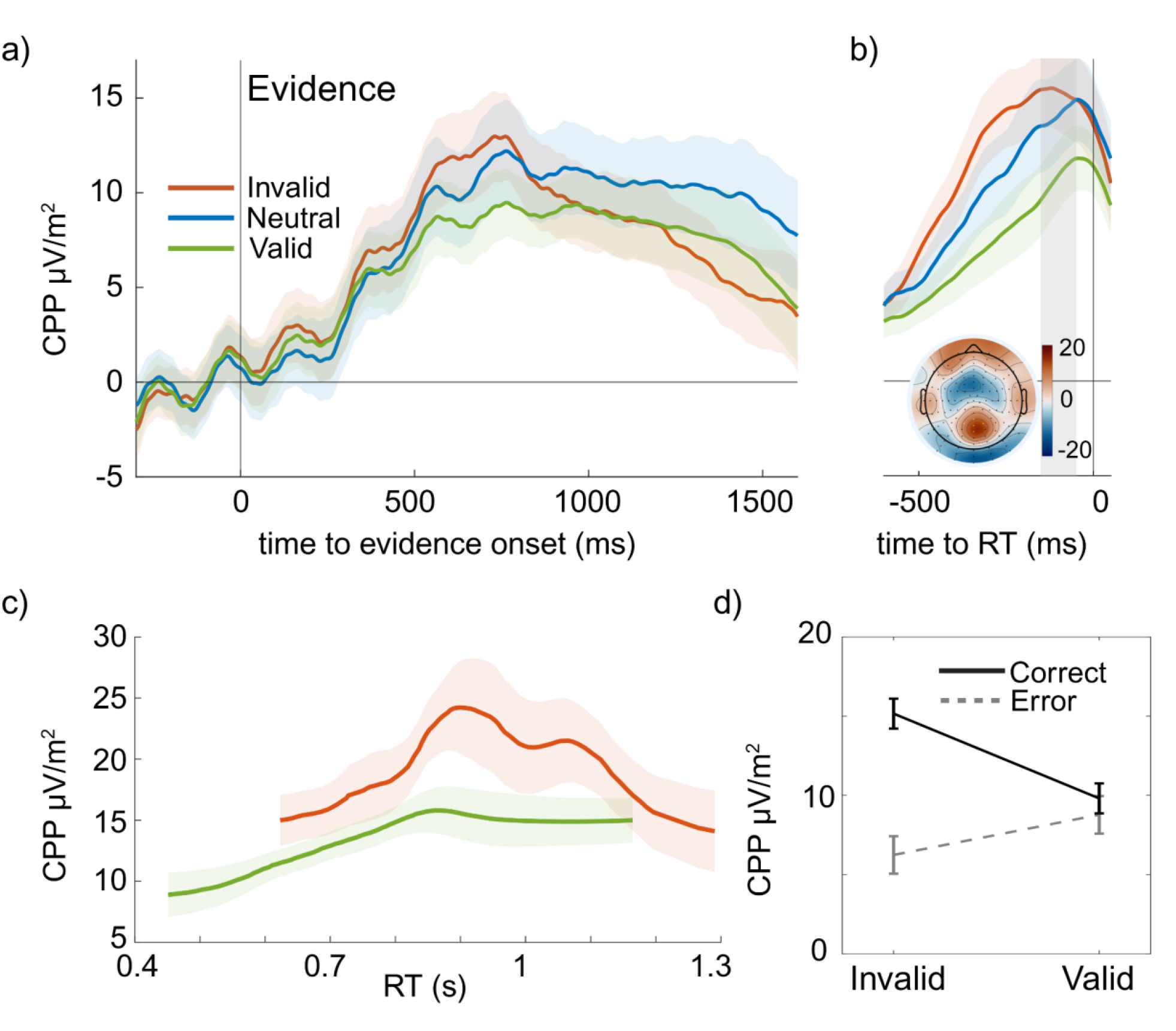
CPP amplitude after matching cue conditions for accuracy (correct trials only) and confidence (‘certain’ ratings only). (a) Stimulus-locked CPP and (b) response-locked CPP exhibit lower amplitudes on valid compared to invalid and neutral cues (n = 19). Grey area (b) shows window used to calculate mean CPP amplitudes. Insets plot (b) shows grand average topography for correct, validly cued trials rated as ‘certain’. (c) CPP amplitude prior to response plotted as a function of reaction time (n = 15; four participants were excluded from this figure due to an insufficient number of trials for either condition). It was consistently smaller on valid compared to invalid cue trials irrespective of RT. (d) Mean pre-response CPP amplitude for responses rated as ‘certain’ was greater for correct than incorrect responses, with amplitudes decreasing from invalid to valid trials for correct responses, whereas the opposite pattern was observed for incorrect responses (n = 17).

These findings do not contradict previous studies reporting a close relationship between CPP amplitude and subjective confidence when prior knowledge was not available (Dou et al., 2024; Feuerriegel et al., 2022; Gherman & Philiastides, 2015; Grogan et al., 2026). Here, within each cue condition, higher confidence ratings were indeed associated with larger CPP amplitudes. These patterns are consistent with the CPP reflecting a sensory evidence accumulation process that informs confidence judgments, rather than directly encoding confidence itself (Fetsch et al., 2014; Grogan et al., 2026). The fact that participants reported higher confidence on validly cued trials despite having accrued smaller quantities of sensory evidence suggests that their confidence ratings reflected a combination of the prior and the observed. Previous behavioural modelling work has demonstrated that human decision-making strategies are informed by two distinct sources of information: true confidence incorporating prior knowledge and ‘counterfactual confidence’ which represents the strength of evidence for the choice discounting the prior (Zylberberg et al, 2018). Our results point to the possibility that the CPP may trace the latter information source. Consistent with this, Desender et al (2025) identified a candidate frontal signal that was sensitive to both prior beliefs and confidence, and which may serve the role of integrating accumulated evidence with prior information to compute ‘true’ confidence.

How the CPP is generated in the brain remains a mystery. While intracranial EEG studies have identified several brain regions exhibiting similar movement-independent evidence accumulation dynamics (e.g. Gherman et al 2024), the underlying circuitry needed to produce such a signal has not been established. A recent study has demonstrated that, at least in multisensory target detection tasks, the CPP appears to arise from the summation of distinct neural populations coding for auditory and visual targets (Egan et al 2025). Ultimately, recordings from individual neurons within candidate CPP generating regions may be required in order to understand how this signal is generated.

In conclusion, the present findings support the view that the CPP reflects sensory evidence accumulation, while explicit confidence reports reflect the combination of accumulated evidence and prior knowledge. In the absence of predictive cues, the CPP’s amplitude does scale with confidence, as both are driven by the same accumulation process.

## Methods

### Participants

We recruited 20 participants (12 females, age range: 18–26 years) for our study. All participants reported having normal or corrected-to-normal vision and no history of psychiatric disorders, epilepsy, or sensitivity to flashing lights. One participant was excluded from analyses following artefact rejection due to low trial numbers. Participants completed four sessions: one practice session and three EEG sessions on separate days. While the sample size was relatively small, a total of 2592 trials per participant were collected to maximise statistical power. Informed consent was obtained from all participants prior to testing. Participants were compensated €70 for their time, in addition to a performance-based bonus (up to €20) and a bonus for completing all four sessions (€10). The study was approved by the Trinity College Dublin Ethics Committee and conducted in accordance with the Declaration of Helsinki.

### Set-up

The experiment was conducted in a dark, sound-attenuated, and electromagnetically shielded booth to minimise external noise. Participants were seated 60 cm away from the CRT monitor (40.5 cm, 1024×768 resolution, 100 Hz refresh rate). To prevent head movement during the task, participants were asked to place their head on the chin rest. The task was programmed using MATLAB (2013b), and participants were using a keyboard to respond. Eye movements were recorded using the EyeLink 1000 eye tracker (1000 Hz sample rate).

### Contrast Discrimination Task

Participants performed a task in which they were required to report which of two overlaid gratings had the higher contrast. Each trial began with a central yellow fixation square that remained on the screen throughout the whole trial. After 400ms, a predictive cue and two overlaid gratings tilted 45° from the vertical axis (135° for left-tilted and 45° for right-tilted) appeared simultaneously.

The predictive cue, displayed as two arrows on either side of the fixation square, was shown for 300 ms. There were two types of cues: neutral cues for which both arrows were coloured yellow and informative cues which featured one yellow and one blue arrow, with the yellow arrow correctly predicting the target grating (i.e. the one that would have higher contrast) with 80% accuracy.

The gratings were faded-in from 0% to 50% contrast for 400ms, followed by an 800ms lead-in period during which both gratings were maintained at 50% contrast to allow sensory-evoked potentials elicited by the stimuli to resolve prior to decision formation. Throughout their presentation, one of the gratings flickered at 20 Hz, while the other flickered at 25 Hz with the flicker rate of each grating randomly varying across trials. After the lead-in period one grating would increase in contrast while another one would decrease by the same amount with this contrast difference maintained for 2 seconds. Participants reported their grating choice and their confidence in that choice simultaneously by pressing “S”, “D” or “F” key with their left hand to respond, “certain left”, “probably left”, “maybe left” and “J”, “K” or “L” key with their right hand to report “maybe right”, “probably right” or “certain right” respectively. Participants were required to make their response within 1.8 seconds of the contrast change. Participants were provided with feedback on the accuracy of their choices on each trial via a change in the fixation square with green indicating a correct response and red indicating that the participant had either responded too soon, incorrectly or too late. For “Too Early” or “Too Late” responses, the red square was accompanied by the corresponding labels above it. At the same time, participants could get different number of points for their answer depending on their accuracy and confidence level (100 points for “certain” and correct, 96 for “probably” and correct, 84 for “maybe” and correct, 64 for “maybe” and incorrect, 36 for “probably” and incorrect, 0 for “certain” and incorrect, -100 for too early or too fast responses). This was not displayed on screen on each trial, but the total they had won was shown after each block and affected their performance bonus payout.

### Training

Each participant underwent a six-block training session. The first five blocks used only neutral cues, with predictive cues introduced in the last block. First, participants completed 50 practice trials to familiarise themselves with the task and response mechanism. The gratings initially had contrasts of 100% and 0%, and through the staircase procedure the contrast difference was decreased after two consecutive correct responses and increased after two consecutive errors. A preliminary difficulty level was established at the end of this block.

Next, two blocks (50 and 60 trials) were used to estimate and correct for the perceptual biases associated with the different grating flicker rates, as higher flicker frequencies can reduce perceived contrast (Devine et al., 2019, Walsh et al., 2024). The first used a staircase procedure to estimate the contrast adjustments required to compensate for this bias. The second used a one-up-one-down staircase in which 25Hz grating was always a correct answer and initially received the contrast adjustment from the previous block. Both gratings started at 50% contrast, and the staircase was adjusted until participants reached 50% accuracy, showing that the two gratings were perceptually matched. A 60-trial QUEST block was used to estimate the task difficulty to achieve ∼70% choice accuracy on neutral trials (Watson & Pelli, 1983), followed by 30 verification trials to confirm the flicker-bias and difficulty estimates. If the participants’ accuracy was low on any of these five blocks, they repeated the initial practice block to ensure understanding of the task.

Finally, predictive cues were introduced in the last block of 48 trials. If accuracy for any cue type was lower than 50%, the block was repeated. During the practice blocks, participants received feedback on their performance that included the points they earned for each response. However, during the actual task sessions, points were not displayed, but participants could see their accuracy and total amount of points after the end of each block.

### Procedure

After completing the training session, participants underwent three EEG sessions, each lasting approximately two hours. Each session began with short training blocks to recalibrate the flicker bias and contrast differences.

Across the three sessions, participants completed a total of 18 blocks, with each block consisting of 144 trials. Accuracy on neutral trials was monitored throughout to ensure it remained within the target range of 60% to 80%. If accuracy deviated outside this range, the contrast level was adjusted by ±2% to maintain the desired task difficulty. Within each block, the flicker frequency, the side of the targeted contrast change, and the cue type for each trial were randomised and counterbalanced to minimise potential bias towards the task.

### EEG acquisition and pre-processing

EEG data were recorded using a BioSemi ActiveTwo system with 128 electrodes at a sampling rate of 1024 Hz. Eye movements were monitored using eye-tracker and vertical electrooculogram (VEOG) electrodes positioned above and below the left eye. Preprocessing and analysis of the data were performed using custom MATLAB scripts incorporating functions from the EEGLAB toolbox with ERPLAB plugins (Delorme & Makeig, 2004; Lopez-Calderon & Luck, 2014).

The preprocessing pipeline included linear detrending to reduce slow drifts and low-pass filtering with a cutoff frequency of 40 Hz (FIR) to remove high-frequency noise. Data were segmented into epochs ranging from 200 ms before cue onset to 2100 ms after evidence onset. Epochs were baseline-corrected using the 200 ms interval prior to cue onset, as this was the latest common period across all cue conditions. Response-locked epochs were extracted from the evidence-locked data, extending from −1600 to 250 ms relative to the response, and baseline-corrected using the mean amplitude in the −300 to 0 ms window before evidence onset.

Channels with extreme variance were interpolated. Epochs were excluded if scalp electrode voltages exceeded 100 µV or if the eye tracker registered blinks or saccades larger than 2° off the fixation square. Participants were excluded from the analysis if fewer than 50% of their trials remained after artefact rejection (1 participant). The Current Source Density (CSD) transformation (Kayser & Tenke, 2015) was applied to the epochs to reduce the spatial overlap of EEG components.

### Analysis

Trials with RTs below 100 ms from stimuli onset or missing responses were excluded from the analysis. Additionally, a minimum of 10 trials was required for each combination of confidence (*maybe, probably, certain*), accuracy (*correct, incorrect*), and/or cue type (*neutral, invalid, valid*) to ensure reliable analysis. For participants whose data did not meet this criterion in certain combinations, those specific trials were excluded, while their remaining data was kept for further analysis (Grogan et al., 2023).

The primary neural signals of interest were the MB rhythm and the CPP. To identify suitable electrodes for representing the CPP, we plotted the grand-average ERP topography across all trials within the time range from -150 to -50 ms relative to response. Based on this topography, we averaged across five electrodes located around standard sites CPz and Pz (ActiCap, Brain Products) to obtain the CPP measurements. Mean CPP amplitudes were calculated within -300:0 ms the window relative to evidence onset for evidence-locked analysis, and within -150:-50 ms window before response for response-locked CPP. For the MB signal analysis, we applied the Short Time Fourier Transform (STFT) to extract activity in the 8–30 Hz frequency range, using a 400 ms window and step size 50 ms. Clusters of three electrodes in each hemisphere were selected for MB analysis, centred on the regions showing the strongest ipsilateral minus contralateral pre-response amplitude (-150 to -50 ms to response) at the grand-average topography and centred around the C3 and C4.

We analysed trial-wise data using mixed-effects models (LMEs), with all continuous predictors z-scored. Binary outcomes such as accuracy were modelled using generalised LMMs with a logistic link. All models included a random intercept for participants to account for individual variability (Smith, P.L. & Little, D.R., 2018).

## Data and code availability

Task scripts and MATLAB code for analysing the data are available at https://codeberg.org/monakhvd/PriorConfidence and anonymous data are available at 10.5281/zenodo.23598820.

## Acknowledgements

This study was supported by a grant from Horizon 2020 European Research Council Consolidator Grant IndDecision 865474.

## References

Carandini, M., & Churchland, A. K. (2013). Probing perceptual decisions in rodents. Nature Neuroscience, 16, 824–831. 10.1038/nn.3410

de Lange, F. P., Rahnev, D. A., Donner, T. H., & Lau, H. (2013). Prestimulus oscillatory activity over motor cortex reflects perceptual expectations. Journal of Neuroscience, 33(4), 1400–1410. 10.1523/JNEUROSCI.1094-12.2013

Delorme, A., & Makeig, S. (2004). EEGLAB: An open source toolbox for analysis of single-trial EEG dynamics including independent component analysis. Journal of Neuroscience Methods, 134, 9–21. 10.1016/j.jneumeth.2003.10.009

Desender, K., Smet, A., Erdil, D., Gunay, E. M., Visser, Y. F., Le Denmat, P., & Van Marcke, H. (2025). Common neural choice signals reflect accumulated evidence, not confidence. Cerebral Cortex, 35(8), bhaf257. 10.1093/cercor/bhaf257

Devine, C. A., Gaffney, C., Loughnane, G. M., Kelly, S. P., & O’Connell, R. G. (2019). The role of premature evidence accumulation in making difficult perceptual decisions under temporal uncertainty. eLife, 8:e48526. 10.7554/eLife.48526

Donner, T. H., Siegel, M., Fries, P., & Engel, A. K. (2009). Buildup of choice-predictive activity in human motor cortex during perceptual decision making. Current Biology, 19(18), 1581–1585. 10.1016/j.cub.2009.07.066

Dou, W., Martinez Arango, L. J., Castaneda, O. G., Arellano, L., Mcintyre, E., Yballa, C., & Samaha, J. (2024). Neural signatures of evidence accumulation encode subjective perceptual confidence independent of performance. Psychological Science, 35(7), 760–779. 10.1177/09567976241246561

Egan, J. M., Gomez-Ramirez, M., Foxe, J. J., O’Connell, R. G., & Kelly, S. P. (2026). Distinct audio and visual accumulators co-activate motor preparation for multisensory detection. Nature Human Behaviour, 10(1), 49–63. 10.1038/s41562-025-02280-9

Fetsch, C. R., Kiani, R., & Shadlen, M. N. (2014). Predicting the accuracy of a decision: A neural mechanism of confidence. Cold Spring Harbor Symposia on Quantitative Biology, 79, 185–197. 10.1101/sqb.2014.79.024893

Feuerriegel, D., Murphy, M., Konski, A., Mepani, V., Sun, J., Hester, R., & Bode, S. (2022). Electrophysiological correlates of confidence differ across correct and erroneous perceptual decisions. NeuroImage, 259, Article 119447. 10.1016/j.neuroimage.2022.119447

Gherman, S., Markowitz, N., Tostaeva, G., Espinal, E., Mehta, A. D., O’Connell, R. G., Kelly, S. P., & Bickel, S. (2024). Intracranial electroencephalography reveals effector-independent evidence accumulation dynamics in multiple human brain regions. Nature Human Behaviour, 8(4), 758–770. 10.1038/s41562-024-01824-9

Gherman, S., & Philiastides, M. G. (2015). Neural representations of confidence emerge from the process of decision formation during perceptual choices. NeuroImage, 106, 134–143. 10.1016/j.neuroimage.2014.11.036

Gold, J. I., & Shadlen, M. N. (2007). The neural basis of decision making. Annual Review of Neuroscience, 30, 535–574. 10.1146/annurev.neuro.29.051605.113038

Grogan, J. P., Rys, W., Kelly, S. P., & O’Connell, R. G. (2023). Confidence is predicted by pre- and postchoice decision signal dynamics. Imaging Neuroscience, 1, imag-1-00005. 10.1162/imag_a_00005

Grogan, J. P., Vermeylen, L., Mannion, S. L., McCabe, C., Monakhovych, D., Desender, K., & O’Connell, R. G. (2026). A single evidence accumulation process informs perceptual choices and subsequent confidence reports. eLife, 15:RP112343. 10.7554/eLife.112343.1

Hanks, T. D., & Summerfield, C. (2017). Perceptual decision making in rodents, monkeys, and humans. Neuron, 93, 15–31. 10.1016/j.neuron.2016.12.003

Hebart, M. N., Schriever, Y., Donner, T. H., & Haynes, J. D. (2016). The relationship between perceptual decision variables and confidence in the human brain. Cerebral Cortex, 26(1), 118–130. 10.1093/cercor/bhu181

Herding, J., Ludwig, S., von Lautz, A., Spitzer, B., & Blankenburg, F. (2019). Centro-parietal EEG potentials index subjective evidence and confidence during perceptual decision making. NeuroImage, 201, 116011. 10.1016/j.neuroimage.2019.116011

Kayser, J., & Tenke, C. E. (2015). On the benefits of using surface Laplacian (current source density) methodology in electrophysiology. International Journal of Psychophysiology, 97(3), 171–173. 10.1016/j.ijpsycho.2015.06.001

Kelly, S. P., Corbett, E. A., & O’Connell, R. G. (2021). Neurocomputational mechanisms of prior-informed perceptual decision-making in humans. Nature Human Behaviour, 5, 467–481. 10.1038/s41562-020-00967-9

Kelly, S. P., & O’Connell, R. G. (2013). Internal and external influences on the rate of sensory evidence accumulation in the human brain. Journal of Neuroscience, 33(50), 19434–19441. 10.1523/JNEUROSCI.3355-13.2013

Kiani, R., Corthell, L., & Shadlen, M. N. (2014). Choice certainty is informed by both evidence and decision time. Neuron, 84, 1329–1342. 10.1016/j.neuron.2014.12.015

Kiani, R., Hanks, T. D., & Shadlen, M. N. (2008). Bounded integration in parietal cortex underlies decisions even when viewing duration is dictated by the environment. Journal of Neuroscience, 28(12), 3017–3029. 10.1523/JNEUROSCI.4761-07.2008

Kohl, C., Spieser, L., Forster, B., Bestmann, S., & Yarrow, K. (2020). Centroparietal activity mirrors the decision variable when tracking biased and time-varying sensory evidence. Cognitive Psychology, 122, 101321. 10.1016/j.cogpsych.2020.101321

Lopez-Calderon, J., & Luck, S. J. (2014). ERPLAB: An open-source toolbox for the analysis of event-related potentials. Frontiers in Human Neuroscience, 8, Article 213. 10.3389/fnhum.2014.00213

Loughnane, G. M., Newman, D. P., Bellgrove, M. A., Lalor, E. C., Kelly, S. P., & O’Connell, R. G. (2016). Target selection signals influence perceptual decisions by modulating the onset and rate of evidence accumulation. Current Biology, 26(4), 496–502. 10.1016/j.cub.2015.12.049

Murphy, P. R., Robertson, I. H., Harty, S., & O’Connell, R. G. (2015). Neural evidence accumulation persists after choice to inform metacognitive judgments. eLife, 4:e11946. 10.7554/eLife.11946

O’Connell, R. G., Dockree, P. M., & Kelly, S. P. (2012). A supramodal accumulation-to-bound signal that determines perceptual decisions in humans. Nature Neuroscience, 15, 1729–1735. 10.1038/nn.3248

O’Connell, R. G., & Kelly, S. P. (2021). Neurophysiology of human perceptual decision-making. Annual Review of Neuroscience, 44:495–516. 10.1146/annurev-neuro-092019-100200

Pleskac, T. J., & Busemeyer, J. R. (2010). Two-stage dynamic signal detection: A theory of choice, decision time, and confidence. Psychological Review, 117(3), 864–901. 10.1037/a0019737

Smith, P. L., & Little, D. R. (2018). Small is beautiful: In defense of the small-N design. Psychonomic Bulletin & Review, 25, 2083–2101. 10.3758/s13423-018-1451-8

Steinemann, N. A., O’Connell, R. G., & Kelly, S. P. (2018). Decisions are expedited through multiple neural adjustments spanning the sensorimotor hierarchy. Nature Communications, 9, 3627. 10.1038/s41467-018-06117-0

Tarasi, L., Covelli, M., Tabarelli de Fatis, C., & Romei, V. (2025). Prior information shapes perceptual confidence. Journal of Cognition, 8(1), 11. 10.5334/joc.417

Twomey, D. M., Kelly, S. P., & O’Connell, R. G. (2016). Abstract and effector-selective decision signals exhibit qualitatively distinct dynamics before delayed perceptual reports. Journal of Neuroscience, 36(28), 7346–7352. 10.1523/JNEUROSCI.4162-15.2016

Urai, A. E., & Pfeffer, T. (2014). An action-independent signature of perceptual choice in the human brain. The Journal of Neuroscience, 34(15), 5081–5082. 10.1523/JNEUROSCI.0477-14.2014

Van Marcke, H., Denmat, P. L., Verguts, T., & Desender, K. (2024). Manipulating prior beliefs causally induces under- and overconfidence. Psychological Science, 35(4), 358–375. 10.1177/09567976241231572

van Vugt, M. K., Beulen, M. A., & Taatgen, N. A. (2019). Relation between centro-parietal positivity and diffusion model parameters in both perceptual and memory-based decision making. Brain Research, 1715, 1–12. 10.1016/j.brainres.2019.03.008

Walsh, K., McGovern, D. P., Dully, J., Kelly, S. P., & O’Connell, R. G. (2024). Prior probability cues bias sensory encoding with increasing task exposure. eLife, 12:RP91135. 10.7554/eLife.91135.3

Watson, A. B., & Pelli, D. G. (1983). Quest: A Bayesian adaptive psychometric method. Perception & Psychophysics, 33, 113–120. 10.3758/BF03202828

Yates, J. L., Park, I. M., Katz, L. N., Pillow, J. W., & Huk, A. C. (2017). Functional dissection of signal and noise in MT and LIP during decision-making. Nature Neuroscience, 20, 1285–1292. 10.1038/nn.4611

Zylberberg, A., Wolpert, D. M., & Shadlen, M. N. (2018). Counterfactual reasoning underlies the learning of priors in decision making. Neuron, 99, 1083–1097.e6. 10.1016/j.neuron.2018.07.035

