## Supplemental Figure 1 for "Distinguishing evidence accumulation from choice confidence in neural signatures of perceptual decision making"

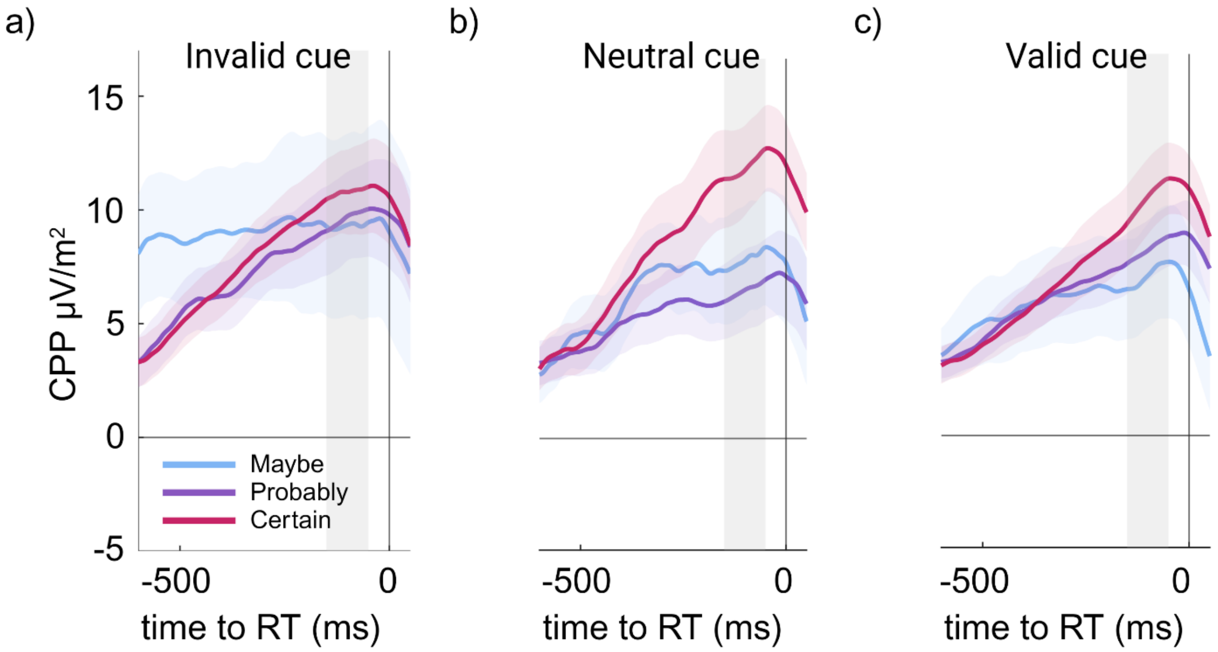


*Supplementary Figure 1. Response-locked CPP waveforms split by confidence ratings ('maybe', 'probably', and 'certain') for (a) invalid, (b) neutral, and (c) valid cues across all trials. Higher subjective confidence levels were associated with higher CPP amplitude across all cue types (n = 19). Grey shading indicates the pre-response time window (−150 to −50 ms) used to calculate mean CPP amplitudes for statistical analyses.*
